# Chemokine CXCL4 interactions with extracellular matrix proteoglycans mediate wide-spread non-receptor mediated immune cell recruitment

**DOI:** 10.1101/2022.07.10.499282

**Authors:** Anna L Gray, Richard Karlsson, Abigail RE Roberts, Amanda JL Ridley, Nabina Pun, Catherine Hughes, Laura Medina-Ruiz, Holly L Birchenough, Catherina L Salanga, Edwin A Yates, Jeremy E Turnbull, Tracy M Handel, Gerard J Graham, Thomas A Jowitt, Ingo Schiessl, Ralf P Richter, Rebecca L Miller, Douglas P Dyer

## Abstract

Leukocyte recruitment from the vasculature into tissues is a crucial component of the immune system, but is also key to inflammatory disease. Chemokines are central to this process but have yet to be therapeutically targeted during inflammation, due to a lack of mechanistic understanding. Specifically, CXCL4 (PF4) has no established receptor that explains its function. Here we use biophysical, *in vitro* and *in vivo* techniques to determine the mechanism underlying CXCL4 mediated leukocyte recruitment. We demonstrate that CXCL4 binds to glycosaminoglycan (GAG) sugars within the endothelial extracellular matrix resulting in increased vascular permeability and non-specific recruitment of a range of leukocytes. Furthermore, GAG sulphation confers selectivity onto chemokine localisation. These findings represent a new understanding of chemokine biology, providing novel mechanisms for future therapeutic targeting.

**One sentence summary:** CXCL4 binds to extracellular matrix proteoglycans resulting in increased vascular permeability and recruitment of a wide range of different leukocytes via a non-canonical mechanism.

## Introduction

Leukocyte recruitment is central to fighting infection, coordinating the immune response to injury, and driving inflammatory disease (*1*). Chemokines are critical during leukocyte recruitment, facilitating firm adhesion of leukocytes to the vascular endothelium (*2*). Due to the central role of chemokines in many diseases, they are prime therapeutic targets (*3*), but drugs that successfully target them are lacking (*4*). This is, in part, due to an under-developed understanding of the mechanisms underlying chemokine biology.

The dogma of the chemokine field states that they bind to G protein-coupled receptors (GPCRs) on leukocytes, producing integrin activation and firm adhesion to the vascular endothelium (*2*). However, this does not hold for CXCL4 (platelet factor 4, PF4) (*5*). CXCL4 is produced by platelets and macrophages, regulates leukocyte recruitment and is implicated in a range of diseases (*6*). However, it does not have a clearly defined receptor that explains its ability to recruit leukocytes, limiting drug discovery efforts (*5*).

Extracellular matrix proteoglycans are present on most cells and are particularly prominent on the vascular endothelium lining blood vessels where they form the endothelial glycocalyx (*7*). This barrier regulates vascular permeability, interactions between circulating leukocytes and the vascular endothelium and is thus a central component of leukocyte recruitment into tissues. Chemokine interactions with key glycocalyx components, glycosaminoglycan (GAG) sugars, are critical for chemokine function *in vivo* but do not directly affect chemokine:receptor interactions (*8*).

The functional role of chemokine:GAG interactions is proposed to be localisation of chemokines to the endothelial surface to ensure that chemokine:receptor interactions occur at a specific site (*9*). This does not explain why chemokines have a wide range of GAG-binding affinities, where some, like CXCL4, have very high-affinity GAG interactions and also crosslink GAG chains (*9–11*). Thus, the biological consequences of CXCL4:GAG interactions have yet to be understood. GAGs and their larger proteoglycan structures can directly mediate cell signalling in other contexts (*12*), but the relevance of this to chemokine function has been largely overlooked. Currently we do not understand the mechanisms that drive CXCL4 mediated leukocyte recruitment and associated disease (*6*). Interactions with extracellular matrix GAGs may provide an “atypical” mechanistic explanation for CXCL4 function.

Herein, we demonstrate that CXCL4 increases endothelial permeability by binding to GAGs rather than chemokine receptors, which results in the recruitment of a wide range of leukocytes. It is the first example of a chemokine functioning primarily through GAGs and not via a receptor. We also show that GAG fine structure (sulphation pattern) confers specificity to chemokine:GAG interactions, which illustrates how GAGs can impose selectivity onto the supposedly “redundant” chemokine system.

## Results

### CXCL4 atypically recruits a broad spectrum of different leukocytes

To understand the mechanism underlying CXCL4 mediated leukocyte recruitment, we used an *in vivo* leukocyte recruitment assay to dissect this process (Fig. 1A). CXCL4 increased the number of CD45^+^ (general leukocyte marker) cells, compared to vehicle controls (Fig. 1B). Surprisingly, CXCL4 produced a significant increase in a wide range of different leukocyte types (Fig. 1C, gating strategy in fig. S1A); specifically, neutrophils, eosinophils, monocytes, dendritic cells, and T cells (TCR beta^+^, NK1.1^+^ and Gamma delta^+^). In contrast, CCL2, which serves as a comparator for CXCL4, given its well established classical recruitment mechanism via leukocyte chemokine receptors (*2*), recruited only CCR2^+^ cells (monocytes, DCs and NK1.1 T cells; fig. S1B).

**Figure 1.**
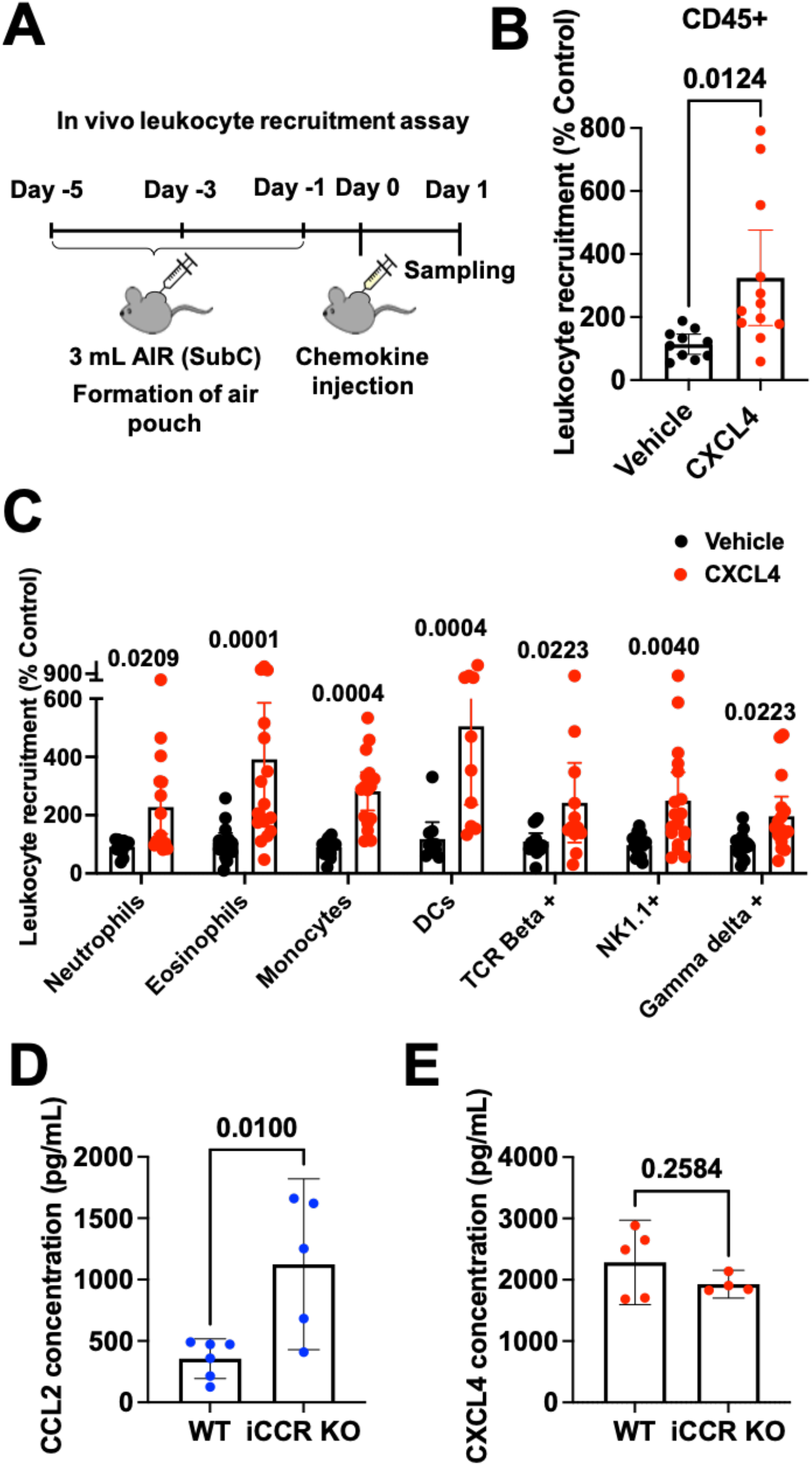
CXCL4 recruits a wide range of different leukocytes *in vivo*. (A) Schematic of the *in vivo* leukocyte recruitment assay. (B) CD45^+^ cell counts 24 hr after CXCL4 injection. (C) Quantification of different leukocytes recruited by CXCL4. (D) CCL2 and (E) CXCL4 quantification in the serum of wild type or iCCR (CCR1, 2, 3 and 5) KO inflamed mice. All plots are mean with 95% confidence intervals and represent at least two separate experiments, data have been pooled and each dot represents an individual mouse. B and C are normalised to vehicle controls. Individual p values are shown, B, D and E analysed using an unpaired t test and C analysed using a one-way ANOVA with a post-hoc Sidak analysis of log-transformed data.

Recruitment of human monocytes by CXCL4 has been reported to be mediated by CCR1 (*13*). However, CXCL4-mediated recruitment of CD45^+^ cells or monocytes remained unchanged in the CCR1 KO mouse compared to wild type controls (fig. S2A). CXCR3 has also been associated with CXCL4 function, albeit at high concentrations (*6*). Analysis of cells recruited by CXCL4 into the air pouch demonstrated that only T cells expressed CXCR3 (fig. S2B). Furthermore, the Immgen database demonstrates that of the leukocytes recruited by CXCL4, only T cells express CXCR3 (*14*). Thus, our analysis showed that there was no chemokine receptor expression pattern across the leukocyte types recruited by CXCL4 that would explain their recruitment. Therefore the mechanism underlying widespread CXCL4-mediated recruitment remained unclear.

We next sought to determine whether leukocytes can directly bind and use CXCL4 to facilitate their migration. We exploited previous observations that when chemokine receptors are knocked out, their ligands are increased in the serum due to reduced uptake and degradation (*15*). For example iCCR (CCR1, 2, 3 and 5 combined) KO mice fail to recruit monocytes to the carrageenan inflamed air pouch (*15*). As a result of the ablated monocyte migration the chemoattractant CCL2 is no longer being used (bound and degraded) during monocyte egress from the blood into the inflamed air pouch of iCCR KO mice, and is found at elevated levels in the serum (Fig. 1D). In contrast CXCL4 levels remained unchanged under the same conditions (Fig. 1C). This suggests that monocytes do not directly utilise CXCL4 during trafficking to the inflamed air pouch, even though CXCL4 facilitated their recruitment. Together these data suggest that CXCL4-mediated recruitment is not driven by binding to chemokine receptors on the leukocyte surface. We hypothesised that CXCL4 would increase the permeability of the vascular endothelium, resulting in a widespread increase of recruited leukocytes.

### CXCL4 increases endothelial permeability

In a Transwell cell migration system (Fig. 2A), purified human monocytes produced a classic chemotaxis response to CCL2 (fig. S3A). Similarly, CCL5 (monocytes), CXCL12 (Jurkats), CCL21 (L1.2 cells) and CXCL8 (neutrophils) mediated recruitment of their concomitant receptor-expressing cells at 10 nM concentrations (Fig. 2A). In contrast, no increase in monocyte recruitment was produced by CXCL4, even at a relatively high concentration of 10 nM, and despite the ability of CXCL4 to mediate monocyte recruitment *in vivo* (Fig. 1).

**Figure 2.**
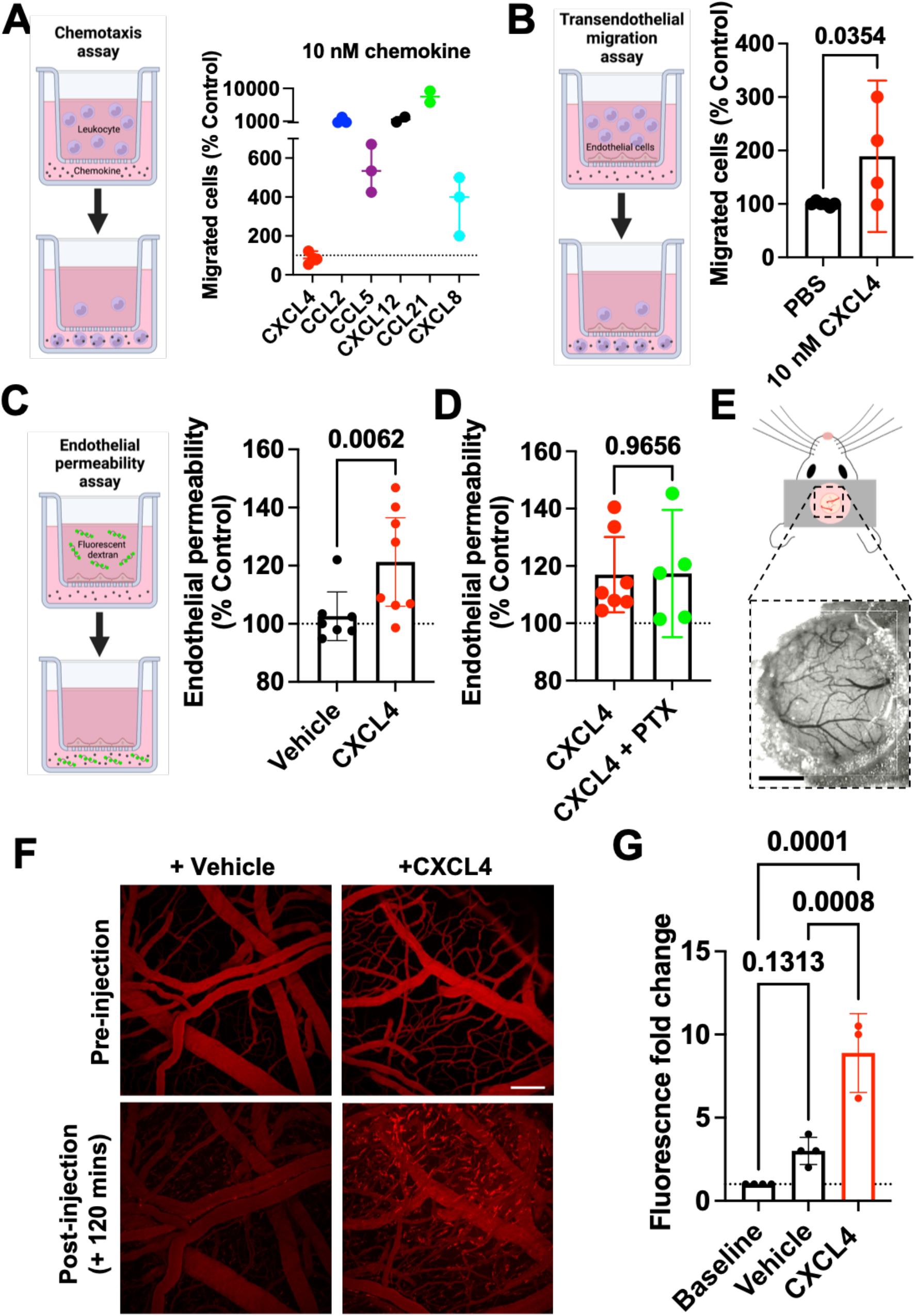
CXCL4 increases endothelial permeability in a receptor independent manner. (A) Chemokine mediated chemotaxis of relevant receptor expressing cells; CXCL4, CCL2 or CCL5 (monocytes), CXCL12 (CXCR4^+^ Jurkat cells), CCL21 (CCR7^+^ L1.2 cells) and CXCL8 (CXCR2^+^ neutrophils). (B) Transendothelial migration of human monocytes towards CXCL4. (C) Transwell endothelial permeability in the absence and presence of CXCL4, (D) CXCL4 alone or in combination with pertussis toxin. (E) Schematic of cranial window implantation for *in vivo* vascular permeability analysis, scale bar = 1mm. (F) *In vivo* analysis of leakage of intravenously injected fluorescent dextran from the vasculature into the meninges following injection of CXCL4 or vehicle controls, scale bar = 100*µ*m. (G) Quantification of (F). All plots are mean with 95% confidence intervals, represent at least two separate experiments where data have been pooled. Data in A-D normalised to vehicle controls. Individual p values are shown, B-D analysed using an unpaired t test and G using a one-way ANOVA with a post-hoc Tukey analysis.

To better replicate the *in vivo* environment we added an endothelial cell monolayer to the upper Transwell chamber to analyse transendothelial migration (Fig. 2B). Addition of CXCL4 (10 nM) produced a significant increase in movement of monocytes to the lower Transwell chamber, demonstrating that CXCL4 acts on the endothelium not the leukocyte.

Next we showed that CXCL4 produced an increase in endothelial permeability, whereas CCL2 did not (Fig. 2C and fig. S3B and C). To determine whether this effect of CXCL4 was mediated by a classical chemokine receptor (Gαi-coupled), we added pertussis toxin (PTx), but observed no effect at concentrations where PTx inhibits GPCRs (Fig. 2D) (*16*). Crucially, CXCL4 increased vascular permeability *in vivo* as shown by leakage of fluorescent dextran from the brain meningeal vasculature after CXCL4 administration (Fig. 2E-G). These findings suggest that CXCL4 may mediate recruitment of immune cells by increasing endothelial permeability via a receptor-independent mechanism.

### CXCL4 function is directly mediated via its interaction with the endothelial GAG heparan sulphate (HS)

Given the independence of CXCL4 mediated leukocyte recruitment from a classic chemokine receptor (Fig. 1 and 2), its high affinity for GAGs and the presence of proteoglycans within the glycocalyx on the blood exposed surface of the endothelium (Fig. 3A) (*7*), we hypothesised that CXCL4 interaction with GAGs on these proteoglycans would directly facilitate leukocyte recruitment. To determine this, we added chemokine to Chinese hamster ovary (CHO) cells with and without genetic KO of the key GAG synthesis gene (B4galt7) (*17*). CXCL4 accumulation was GAG-dependent and much higher than with CCL2 (Fig. 3B and fig. S4A), despite their similar molecular weight and oligomerisation propensity (*10, 18*). HS is thought to be the dominant GAG within the endothelial glycocalyx (*7*). We, therefore, used a biophysical bio-layer interferometry (BLI) approach (Fig. 3C) and determined that the binding of CXCL4 to an HS GAG surrogate, heparin octasaccharide (dp8), was much higher than for CCL2 (Fig. 3D and fig. S4B). This difference in cellular accumulation and GAG binding indicated that CXCL4 function could be directly dependent on its GAG interaction, unlike CCL2.

**Figure 3.**
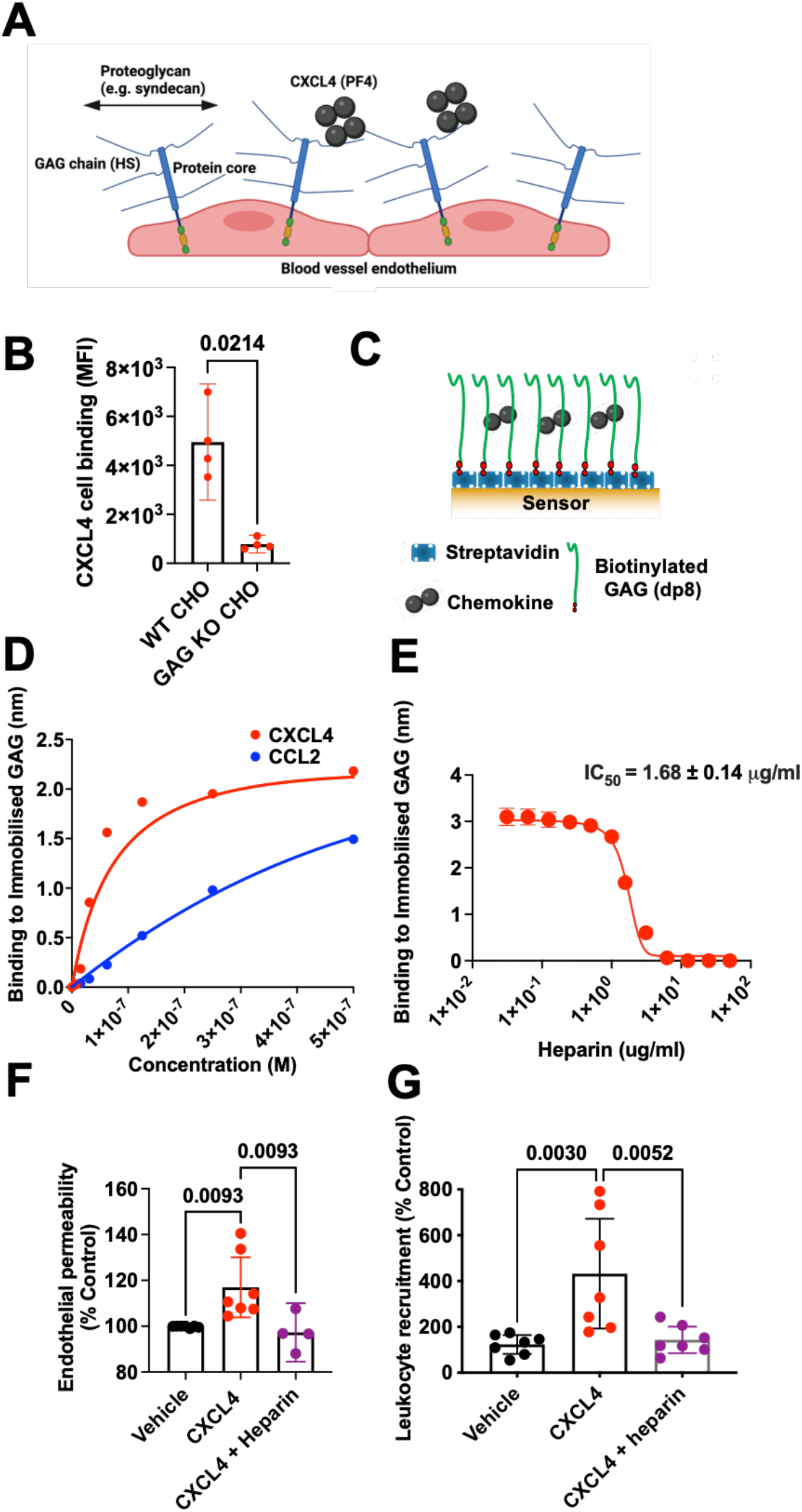
CXCL4 function is mediated by its interaction with glycosaminoglycans (GAGs). (A) Schematic of proteoglycans forming the luminal endothelial glycocalyx. (B) CXCL4 binding to CHO cells with and without surface GAGs. (C) Schematic of the biophysical BLI GAG-binding assay. (D) Maximum signal of chemokine:dp8 binding at different chemokine concentrations in the BLI assay. (E) Maximum CXCL4 (100 nM) binding to dp8 signal following pre-incubation with increasing concentrations of heparin. (F) Endothelial permeability (Transwell) assay with and without CXCL4 and exogenous heparin. (G) *In vivo* leukocyte (CD45^+^) recruitment (air pouch) to CXCL4. B, F and G are mean with 95% confidence intervals and represent at least two separate experiments where data have been pooled and each dot in G represents an individual mouse. D and E are representative of at least two separate experiments. F and G are normalised to vehicle controls. Individual p values are shown, B analysed using an unpaired t test, F and G analysed using a one-way ANOVA with a post-hoc Dunnett analysis of log-transformed data.

To test this hypothesis we firstly showed that CXCL4 binding to GAG was inhibited by pre-incubation with exogenous purified GAG (heparin) in the BLI assay (Fig. 3E and fig. S4C) and on the cell surface (fig. S4D). Subsequently we determined that pre-incubation with heparin reversed CXCL4 mediated increases in endothelial permeability (Fig. 3F) and ablated CXCL4 mediated leukocyte recruitment *in vivo* (Fig. 3G). In contrast, CCL2 mediated recruitment of monocytes to the air pouch was only reduced by a third following pre-incubation with exogenous heparin (fig. S4E).

These experiments demonstrate that the interaction of CXCL4 with GAGs is responsible for CXCL4 mediated recruitment, possibly by producing an increase in endothelial permeability. In contrast the CCL2:GAG interaction has an important but less direct role in facilitating CCL2 mediated leukocyte recruitment; presumably via localisation within the endothelial glycocalyx to facilitate interaction with leukocyte receptors.

### CXCL4 oligomerisation mediates GAG binding and leukocyte recruitment

We have previously shown that high-affinity chemokine:GAG interactions are mediated by chemokine oligomerisation and cross-linking of GAG chains (*10, 11*). Therefore, we next tested the hypothesis that CXCL4 oligomerisation is a critical driver of leukocyte recruitment via crosslinking of GAGs (Fig. 5).

To analyse chemokine oligomerisation we used sedimentation velocity analytical ultra-centrifugation (SV-AUC). SV-AUC showed that CXCL4 has a sedimentation coefficient of 2.7S, with an estimated frictional ratio of 1.33 and a mass estimate using the Svedberg equation of 28.9 kDa, consistent with its formation of tetramers in solution (Fig. 4A) (*19, 20*). The CXCL4 K50E mutant had a much lower sedimentation coefficient of 1.26S, suggestive of either a compact monomer or an elongated dimer; confirming that the K50E mutation inhibits CXCL4 oligomerisation (Fig. 4B) (*20*). CXCL4 K50E also exhibited a large reduction in binding to GAG in the BLI assay, compared to WT CXCL4 (Fig. 4C). We therefore hypothesised that inhibition of CXCL4 oligomerisation is a potential avenue to target CXCL4 driven leukocyte recruitment via inhibition of GAG binding.

**Figure 4.**
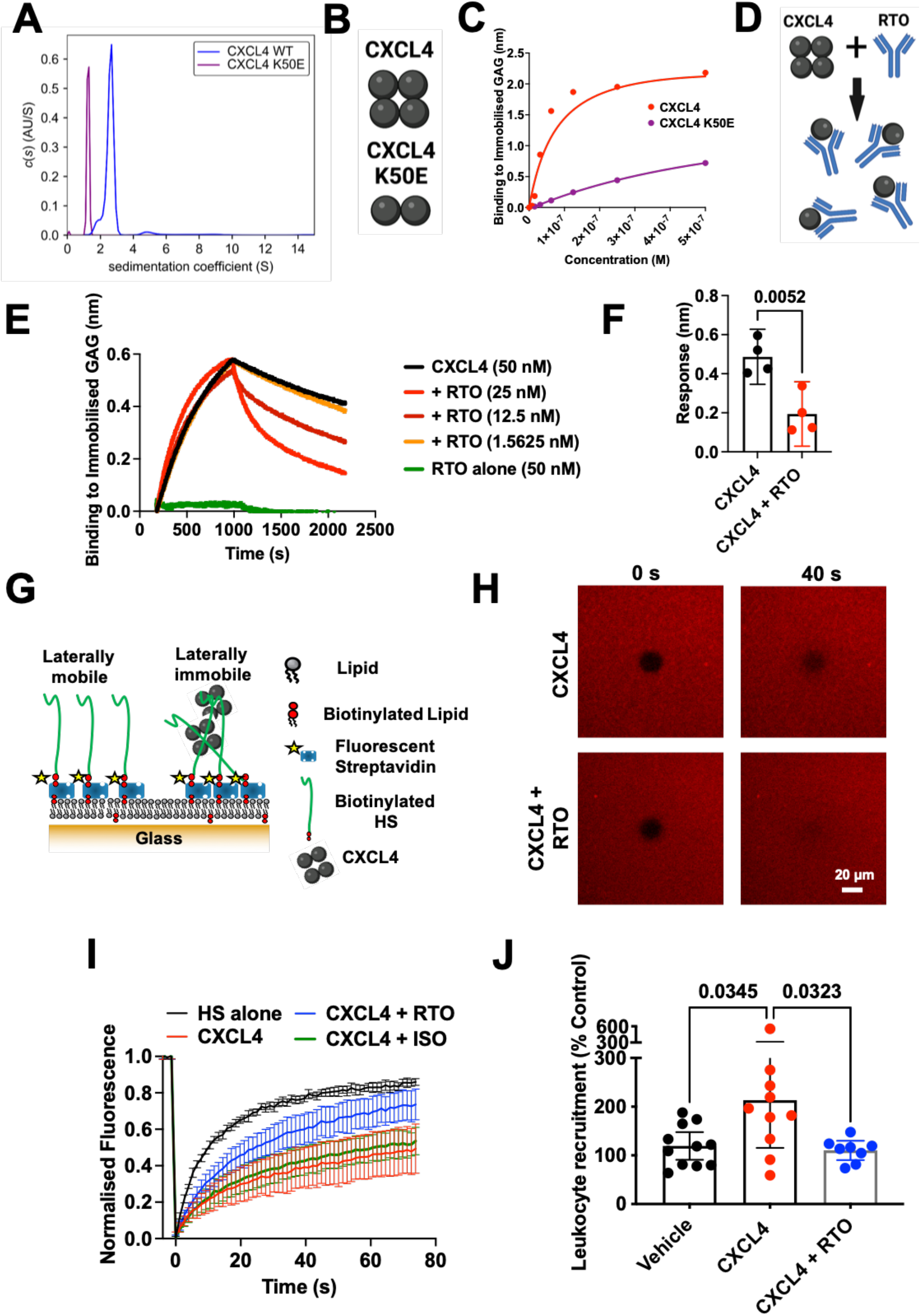
CXCL4 oligomerisation drives leukocyte recruitment. (A) AUC analysis of CXCL4 and the mutant K50E to show sedimentation coefficients (indicative of size). (B) Schematic of oligomerisation states of CXCL4 or the CXCL4 K50E mutant. (C) BLI analysis of CXCL4 K50E binding to immobilised heparin dp8. (D) Schematic of RTO antibody binding to CXCL4 monomer to inhibit oligomerisation. (E) CXCL4 (50 nM) binding to dp8 was monitored in the absence and presence (at a range of concentrations) of RTO antibody. (F) Final CXCL4:dp8 signal (after washing) from (E) is plotted with and without pre-incubation with RTO antibody (25 nM). (G) Schematic of FRAP assay to analyse HS cross-linking. (H) FRAP analysis of GAG in-plane mobility with CXCL4 alone or in combination with the RTO antibody, images post bleaching at indicated times. (I) Fluorescence recovery over time. (J) CXCL4 mediated leukocyte recruitment (CD45^+^) with and without RTO antibody. F, I and J are mean with 95% confidence intervals and represent at least two separate experiments where data have been pooled and each dot in J represents an individual mouse. A, C, E, H and I are representative of at least two separate experiments. Data in J is normalised to vehicle controls. Individual p values are shown, F analysed using an unpaired t test and J analysed using a one-way ANOVA with a post-hoc Dunnett analysis of log-transformed data.

Incubation of CXCL4 with an anti-CXCL4 antibody (RTO) that binds monomeric CXCL4 and inhibits CXCL4 oligomerisation (Fig. 4D) (*21*) produced a dose-dependent inhibition of the CXCL4:GAG interaction in BLI (Fig. 4E-F and fig. S5A). The inhibitory effect of RTO produced an increased off-rate of CXCL4 from GAG, re-creating that observed for CXCL4 K50E (fig. S5B). The inhibitory effect of RTO was specific to CXCL4 since it did not affect CCL2 binding to GAG (fig. S5C) or, bind to the GAG directly (Fig. 4E).

Since the RTO antibody affected the off-rate of the CXCL4:GAG interaction (Fig. 4E), we next sought to determine whether RTO could affect CXCL4 mediated cross-linking of GAG chains. To test this we used a biophysical model of the glycocalyx formed by HS GAG chains anchored on a fluid supported lipid bilayer, enabling analysis of GAG in-plane mobility and cross-linking using fluorescence recovery after photo-bleaching (FRAP) (Fig. 4G-I and fig. S6). CXCL4 inhibited recovery 40 seconds after bleaching (Fig. 4H) and over time (Fig. 4I), indicating cross-linking of GAG chains. Pre-incubation of CXCL4 with RTO returned the bleaching recovery close to that seen in the absence of CXCL4 (HS alone) (Fig. 4H-I and fig. S6). These data demonstrate that RTO mediated inhibition of CXCL4 oligomerisation prevents cross-linking of GAGs. Together, the AUC, BLI and FRAP data demonstrate that chemokine oligomerisation and cross-linking of GAGs are related events that drive the interaction with GAGs.

To determine the importance of CXCL4 oligomerisation, GAG-binding and cross-linking in leukocyte recruitment we utilised RTO in our *in vivo* leukocyte recruitment model (Fig. 4J). RTO resulted in ablation of CXCL4 mediated CD45^+^ leukocyte recruitment, confirming the importance of oligomerisation *in vivo*. This further supports the direct role of GAGs in CXCL4 driven leukocyte recruitment since monomeric chemokines promote chemokine receptor signalling through their GPCRs (*9, 22*). Indeed, recent structural studies suggest that chemokine oligomerisation and receptor binding are likely to be mutually exclusive due to overlap in binding sites, at least for many receptor:chemokine pairs (*8*).

### GAG sulphation confers selectivity on chemokine localisation

The above data show that CXCL4 function is driven by its interaction with endothelial cell surface GAGs. To further characterise chemokine:GAG interactions, we probed the role of specific GAG sulphation positions (Fig. 5A). We hypothesised that changes in GAG sulphation would alter which chemokines are bound and presented on cells.

**Figure 5.**
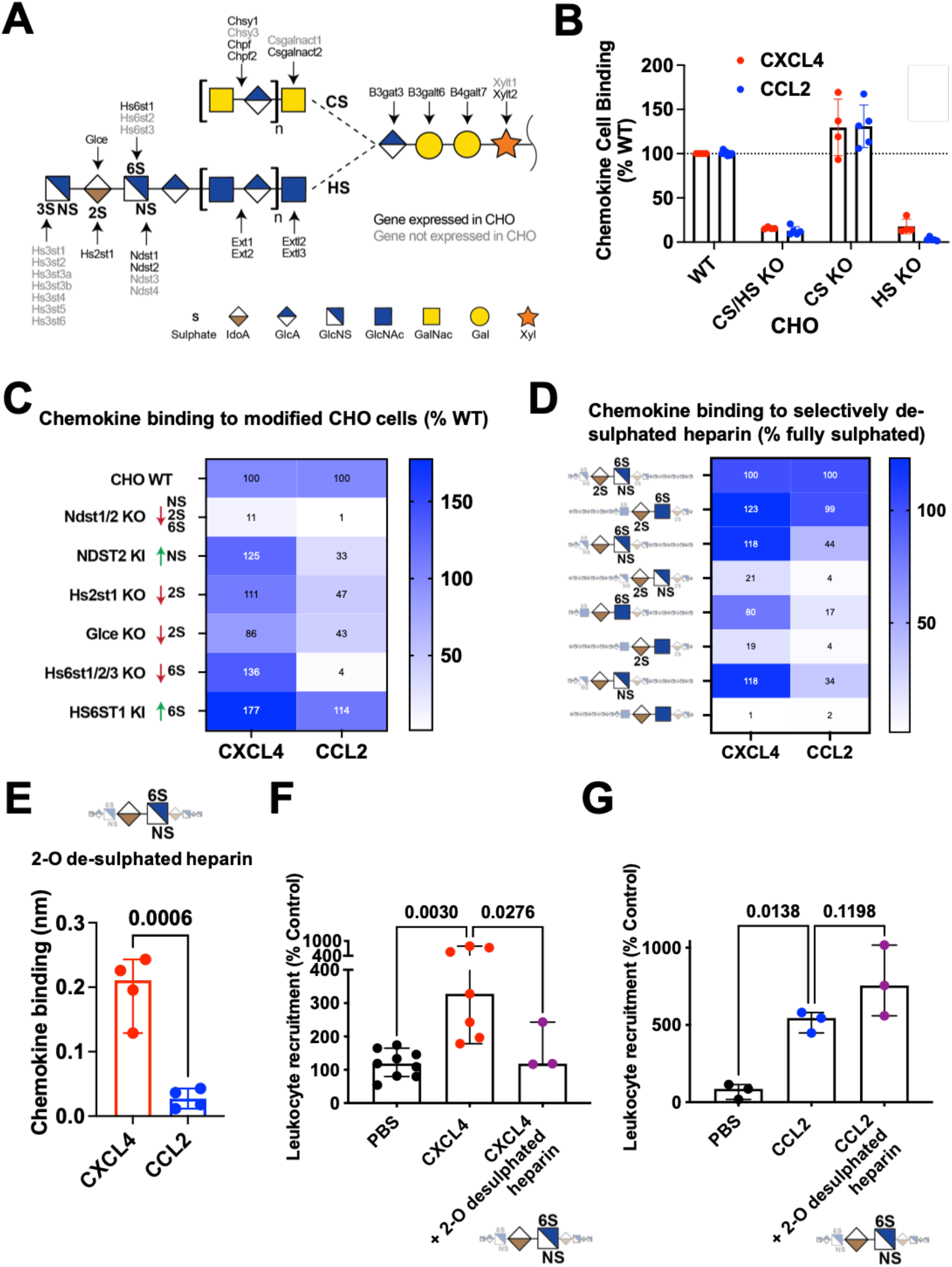
GAG sulphation mediates chemokine interaction selectivity and cellular localisation. (A) Overview of enzymes involved in the biosynthesis of HS and CS GAGs. Elongation proceeds from right to left, a tetrasaccharide linker is elongated with either HS or CS disaccharide repeats into long linear polysaccharides. HS is further modified by addition of sulphate groups at the N-, 2-O, 6-O or 3-O (rarely) positions, this process is catalysed by the indicated sulphotransferases. (B) Chemokine binding to WT CHO cells or CHO cells with no GAGs (KO B4galt7), no CS (KO Csgalnact1/2/Chsy1) or no HS (KO Extl3). (C) Heat map analysis of chemokine binding to CHO cells with KO/KI of sulfotransferases acting on HS, where data is normalised to WT cells. (D) Heatmap analysis of maximum BLI signal of chemokine (500 nM) binding to differentially de-sulphated heparin fragments, data normalised to binding on fully-sulphated heparin. (E) Raw signal of chemokine binding to 2-O de-sulphated heparin in the BLI assay. Chemokine mediated *in vivo* leukocyte recruitment with and without 2-O de-sulphated heparin (F) CD45^+^ cell counts (CXCL4) or (G) monocyte (Ly6C^+^) cell counts (CCL2). B, E, F and G are mean with 95% confidence intervals and represent at least two separate experiments where data have been pooled; each dot in F and G represents an individual mouse. Data in F and G is normalised to vehicle controls. Individual p values are shown, E analysed using an unpaired t test. F and G analysed using a one-way ANOVA with a post-hoc Sidak analysis of log-transformed data.

Earlier experiments (Fig. 3B) demonstrated that KO of CHO GAGs reduced binding to CXCL4. To determine which GAG types are responsible for chemokine binding we used CHO and HEK cells with gene KOs to remove, individually or in combination, HS and/or chondroitin sulphate (CS) (Fig. 5A and fig. S7) (*17, 23*). We determined that HS bound to CCL2 and CXCL4 (Fig. 5B) whilst CS GAG only bound to CXCL4 on HEK cells (fig. S7). HS GAG chains are 20-200 repeating disaccharides in length and are selectively sulphated on the N-, 2-O, 6-O and 3-O positions, which mediate chemokine:GAG binding (Fig. 5A) (*24*). The enzymes that produce these different sulphation points were systematically knocked out in CHO cells, facilitating analysis of their contribution to chemokine:HS interactions (Fig. 5C). KO of NDST1/2, which reduces N-, 2-O and 6-O sulphation on HS (*17*), largely ablated CHO cell binding to CXCL4 and CCL2. CXCL4 binding was not reduced in any of the other cell lines. CCL2 binding was reduced in the absence of HS2ST1 (2-O sulphation), GLCE (required for 2-O sulphation) and combined HS6ST1, 2 and 3 (6-O Sulphation). Since CHO cells do not make 3-O sulphated HS, the genes that produce it were knocked in (*25*). 3-O sulphation generally increased CHO cell interaction with CXCL4 and CCL2 (fig. S7B), however, HS3ST1 KI significantly enhanced binding to CCL2 but not CXCL4. These data demonstrate that differential sulphation of GAG chains provides considerable selectivity in binding to chemokines and presenting them on the cell surface.

### Preferential targeting of CXCL4 function using GAG mimetics

Disruption of chemokine:GAG interactions is a possible avenue for therapeutic intervention in disease (*26*). Heparin GAG and its derivatives are used as anti-coagulants and are well-tolerated in the clinic (*27*). Given the selective interactions mediated by GAG sulphation (Fig. 5A-C), we hypothesised that modified heparin derivatives, with specific sulphation sites removed, may allow preferential inhibition of chemokine driven leukocyte recruitment. To determine binding selectivity of differentially sulphated heparins we used BLI to analyse binding to CXCL4 and CCL2 (Fig. 5D). This confirmed higher accumulation of CXCL4, over CCL2, on fully sulphated (2-O, 6-O and NS) heparin (fig. S7C). Again CCL2 was more sensitive to removal of sulphate groups than CXCL4 (Fig. 5D). The CXCL4 K50E mutant demonstrated that oligomerisation is important for overall GAG binding and not sulphation specificity (fig. S7C and D). These data further demonstrated chemokine:GAG selectivity (*28*).

This BLI approach identified a heparin form (2-O desulphated) with full binding activity for CXCL4 and reduced binding to CCL2 (Fig. 5E). Pre-incubation with the 2-O de-sulphated fragment abolished CXCL4-, but not CCL2-, mediated leukocyte recruitment *in vivo* (Fig. 5F and G). These data demonstrate sulphation-driven selectivity of chemokine:GAG interactions and suggest that this can be exploited to use GAG mimetics to selectively inhibit certain chemokines.

## Discussion

Here we show that the chemokine CXCL4 mediates recruitment of a range of different leukocytes by binding to the endothelial extracellular matrix and not chemokine receptors on leukocytes (Fig. 6). We hypothesise that CXCL4 binding and cross-linking of GAG chains on endothelial proteoglycans triggers signalling mechanisms within the endothelial cells that mediate the increased permeability observed here (Fig. 6) (*12*).

**Figure 6.**
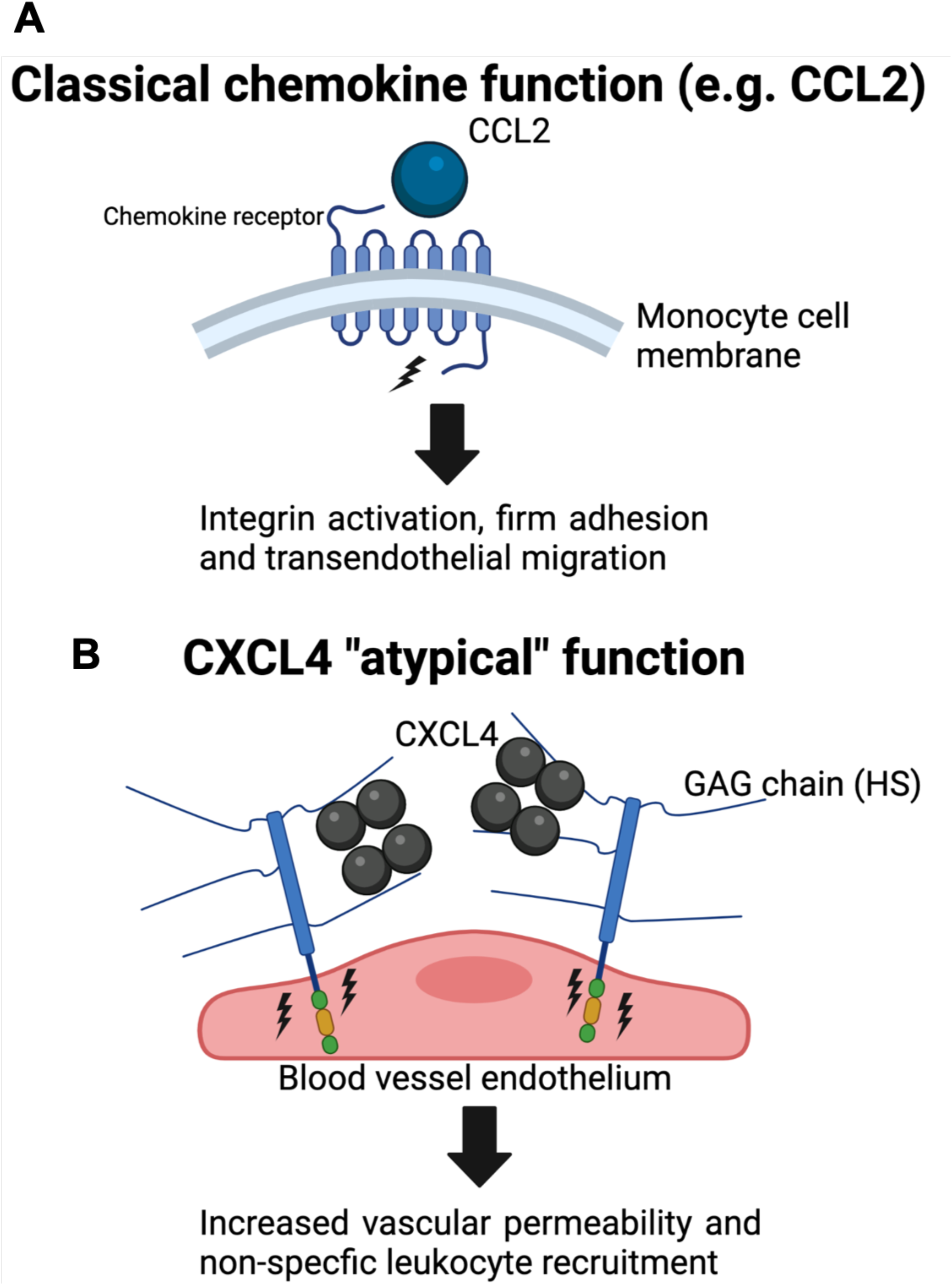
CXCL4 binds to endothelial GAG sugars, resulting in an increase of vascular permeability and non-specific leukocyte recruitment. (A) Classical chemokines, e.g. CCL2, facilitate leukocyte recruitment by binding to seven transmembrane receptors on circulating leukocytes leading to signalling, integrin activation and firm adhesion of the leukocyte to the endothelium. (B) We propose that CXCL4 binds and re-models (as a tetramer) endothelial GAG sugars within the glycocalyx. This produces increased vascular permeability, possibly via signalling through the proteoglycan, facilitating “non-specific” recruitment of a range of different leukocytes from the vasculature and into inflamed tissues.

This generalised recruitment of different leukocytes is supported by previous observations that CXCL4 recruits neutrophils and monocytes in response to ischaemia reperfusion injury (*29*), neutrophils to influenza or *P. aeruginosa* infected lungs (*30, 31*), monocytes to atherosclerotic plaques (*32*) and T cells into the malaria infected brain (*33*). Our study may also provide a mechanism explaining previous observations that CXCL4 increases vascular permeability in the brain during cerebral malaria and in the lung following acute injury (*33, 34*). CXCL4 mediated increase of vascular permeability also fits with the role of platelets in facilitating rapid recruitment of leukocytes in response to inflammation (*35*).

The chemokine system has long been described as redundant, whereby multiple ligands can bind to the same receptor and multiple receptors can bind to the same ligand. However, studies designed to address whether chemokine redundancy exists have found limited evidence for it and in fact suggest specificity (*15, 36–38*). Specificity of receptor function is largely explained by co-ordinated cellular expression (*39*). Contrastingly, there are many examples where chemokine ligands that recruit the same cell types are present at comparable concentrations in the same context (*28*). Our data suggest that chemokine:GAG interactions, co-ordinated by the differential GAG sulphation patterns found across cells and tissues(*40*), will produce specific localisation of chemokines at the cellular and tissue level.

This study demonstrates new ways to target CXCL4 in inflammatory disease by inhibiting oligomerisation and GAG binding using GAG mimetics or specific antibodies. Blocking CXCL4-mediated recruitment of a range of leukocytes and increased vascular permeability is particularly promising in acute hyper-inflamed disease, e.g. sepsis (*34*). Such approaches may also be relevant to targeting CXCL4 in the rare side effects of adenovirus vaccines against SARS-CoV-2 infection (*41*). Future work will be important to address the potential role of CXCL4 as a master regulator of leukocyte recruitment in health and disease.

## Materials and methods

### Materials

Mice were housed in cages of up to four on a 12-h light/dark cycle, with ad libitum access to food and water. All experiments were carried out following ethical approval from The University of Manchester and University of Glasgow and under licence from the UK Home Office (Scientific Procedures Act 1986).

All chemokines were purchased from Protein Foundry. The dp8 and heparin GAGs were purchased (Iduron) and the de-sulphated heparin fractions were generated as detailed in (*42*).

#### Antibodies

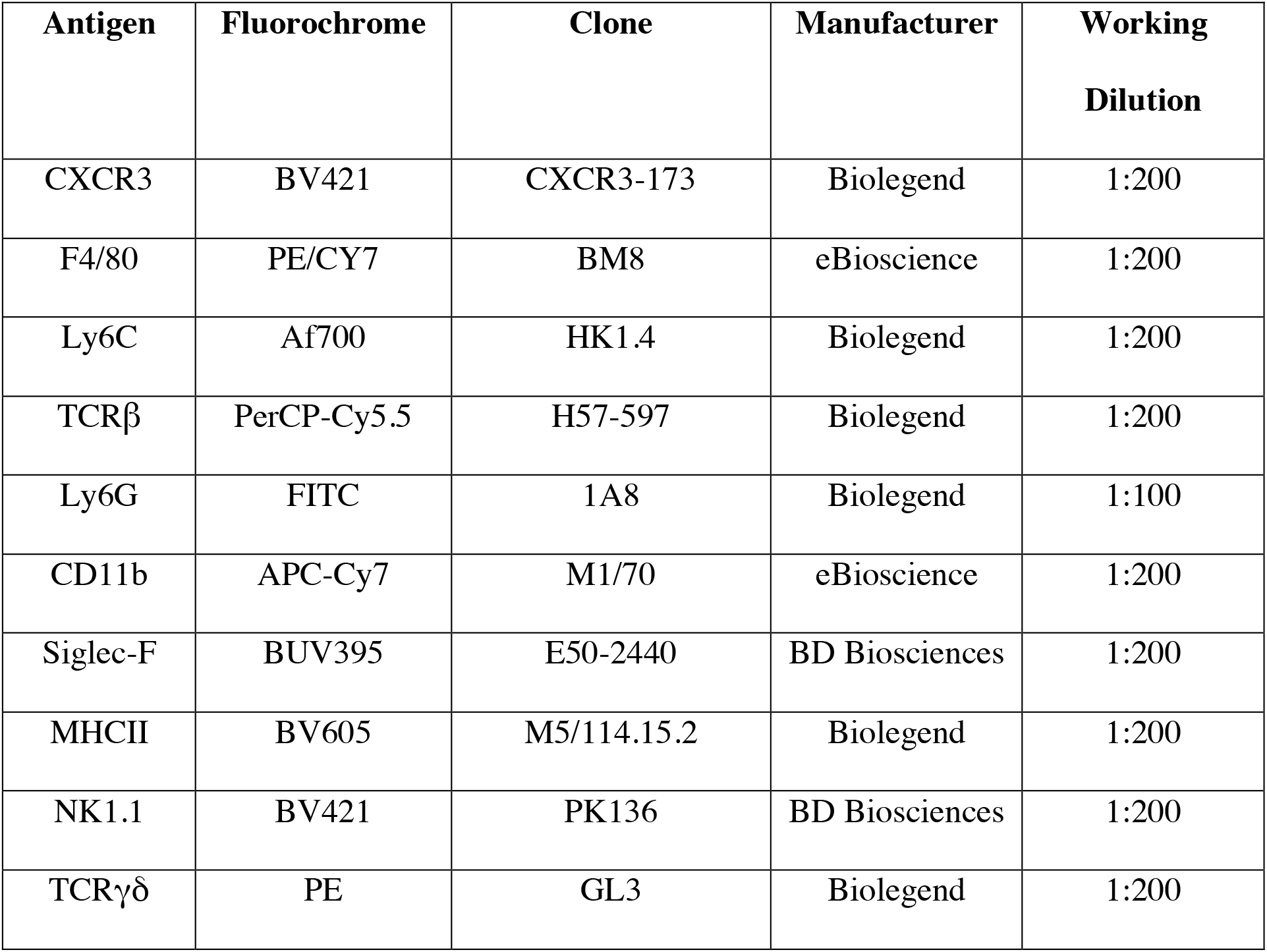

### Methods

#### *In vivo* leukocyte recruitment

The air pouch was created by injecting 3 ml of sterile air subcutaneously under the dorsal skin on 3 occasions 48 hours apart as described previously (*15*). The indicated quantity of chemokine was re-suspended in 100 *µ*l of phosphate buffered saline (PBS (Sigma)) and injected into the air pouch. Mice were culled 24 hours later and the air pouch flushed with 5 ml of flow cytometry buffer (PBS containing 1 mM EDTA (Sigma) and 1% (w/v) foetal bovine serum (Sigma)) to retrieve recruited cells. Retrieved cells were washed in PBS and stained with fixable cell viability dye (eBioscience, 1:1000 in PBS) for 20 minutes at 4 °C before being washed in flow cytometry buffer and re-suspended in 100 *µ*l antibody staining cocktail (antibodies and FcR blocking reagent (Miltenyi Biotec) diluted as indicated above). Cells were incubated for 20 minutes at 4 °C to stain before being washed in flow cytometry buffer and fixed for 20 minutes using 100 *µ*l fixation buffer (BioLegend). CXCR3 staining was performed at 37 °C for 15 minutes prior to surface antibody staining. Cells were then re-suspended in flow cytometry buffer before addition of counting beads (ThermoFisher Scientific) and analysed using a Fortessa flow cytometer (BD Biosciences).

Flow cytometry data was analysed to quantify absolute cell counts or as the percentage of live cells before being normalised relative to vehicle controls to facilitate comparison across experiments.

#### ELISA analysis of CXCL4 and CCL2 concentrations

Specific concentrations of CXCL4 and CCL2 were measured by enzyme-linked immunosorbent assay (ELISA), using Mouse CXCL4/PF4 and Mouse CCL2/JE/MCP-1 DuoSet ELISA (R&D Systems) in a 96-well high binding ELISA plate coated with capture antibody (2 *µ*g/mL), sealed, and incubated overnight at room temperature. The plate was then aspirated and washed with wash buffer composed of 0.05% Tween 20 (Sigma), repeating the process two times for a total of three washes. The remaining wash buffer was removed from the plate by inverting the plate and blotting it against clean paper towels. 150 *µ*l of 1% (w/v) bovine serum albumin (BSA, Sigma) in PBS (Sigma) was used to block the plate and then left to incubate at room temperature for 2 hours. Aspiration and wash steps were repeated, and the plates were ready for sample addition. The standard curve was made using two-fold dilutions and 1% (w/v) BSA was added as a blank. 50 *µ*L of sample or standards in 1% BSA was added to each appropriate well, covered and left to incubate at 4°C overnight. Aspiration was repeated and 50 *µ*L of streptavidin-HRP was added to each well and left to incubate at room temperature for 20 minutes. A thorough aspiration and wash steps were repeated and 50 *µ*L of substrate solution was added to each well and incubated for 20 minutes at room temperature, avoiding exposure to direct light. 25 *µ*L of stop solution was then added to all wells. The optical density of each sample was read out on a Versamax Microplate Reader (Marshall Scientific) at 450nm wavelength.

#### *In vitro* chemotaxis, endothelial transmigration and endothelial permeability Chemotaxis

Monocyte and neutrophil purification was performed as described previously (*18*), briefly blood was obtained from the San Diego Blood Banks and peripheral blood mononuclear cells (PBMCs) were separated by using a Ficoll-Paque gradient and centrifugation. Monocytes were then purified from freshly isolated PBMCs using a CD14^+^ selection MACS purification (Miltenyi Biotec) followed by final re-suspension in chemotaxis buffer (Dulbecco’s modified Eagles medium (DMEM) (Sigma) + 10% (v/v) foetal bovine serum (FBS) (Sigma)). Neutrophils were removed from the bottom of the Ficoll-Pague gradient and red blood cells were removed using red blood cell lysis buffer followed by washing and final re-suspension in chemotaxis buffer.

Chemokine or vehicle control was added in the bottom well of endothelial transwells (5 *µ*m pores (Corning)) at the indicated concentration in 600 *µ*l of chemotaxis buffer. 100 *µ*l of 2 x 10^6^ (2 x 10^4^ total) cell suspension (monocytes (CCL2 and CCL5)), Jurkats (CXCL12), CCR7^+^ L1.2 cells (CCL21) and neutrophils (CXCL8)) were added to the top well for the indicated chemokine. Transwells were then incubated for 2 hours at 37 °C/5 % CO_2,_ removed and faux adherent cells from the lower membrane washed into the bottom well before migrated cells were counted using a Guava EasyCyte 8HT Flow cytometer (Merck Millipore). Data was then presented as the total number of migrated cells or relative to vehicle controls (% control).

##### Transendothelial migration and permeability

Transendothelial migration experiments were undertaken as above but prior to the addition of migrating cells, endothelial cells were coated onto the upper side of the transwell insert as follows. Transwells with 5 *µ*m pores (Corning) were coated with 50 *µ*l per well of 10 *µ*g/ml of fibronectin (Sigma) and incubated for 1 hour at 37 °C/5% CO_2_ and washed three times with 50 *µ*l per well of PBS (Sigma). Immortalised human umbilical vein endothelial cells (Ea.Hy 926 cells) (ATCC) were seeded into each well, 100,000 cells per well in 50 *µ*l of culture medium (DMEM (Sigma) containing 1% (v/v) penicillin-streptomycin solution (Thermo Fisher Scientific), 10% (v/v) FBS (Sigma) and 1% (v/v) L-Glutamine (Sigma)) and incubated for 2 days at 37 °C/5% CO_2_. Transwells were then washed with PBS (three times with 50 *µ*l) and finally chemotaxis buffer prior to endothelial transmigration experiments. In the case of endothelial permeability experiments, 100 *µ*l of chemotaxis buffer containing 25 *µ*M 70,000 MW Texas Red dextran (Thermo Fisher Scientific) was added to the upper side of the endothelial monolayer, following incubation (2 hrs 37 °C/5% CO_2_) fluorescence of the bottom well was determined using a FlexStation 3 fluorescent plate reader (Molecular Devices). Data were expressed and plotted relative to vehicle controls.

#### *In vivo* vascular permeability

##### Animals

Cranial window implantation surgery was performed on male C57BL/6J mice at 20-25 grams and multiphoton imaging was conducted two weeks later.

##### Cranial window implantation

Cranial window implantation was conducted as previously described (*43*). In brief, animals were anaesthetised with 2.5% isoflurane in room air and positioned within a stereotactic frame. The cranium was exposed by removal of the scalp on top of both brain hemispheres. A metal head plate (Narishige CP-2, Japan) was mounted using dental cement (Sun Dental, Japan), to allow later fixation within the multiphoton microscope setup. A 3 mm diameter circular piece of bone was then carefully removed using a dental drill and a circular coverslip (Warner Instruments, USA) was secured in place of the removed bone using dental cement.

##### Multiphoton imaging

Animals underwent intravenous injection of 50 *µ*l of 10 mg/ml neutral 70,000 MW Texas Red dextran (Thermo Fisher Scientific) in sterile saline following induction of anaesthesia. Depth of anaesthesia was maintained throughout the experiment using 1.5-2% isoflurane in 100% oxygen and a heating blanket with temperature probe was used to ensure maintenance of body temperature at 37.5 °C. Images were collected on a Leica SP8 Upright Multiphoton microscope using a Leica 25x / 0.95 L HC Fluotar dipping objective. Images were captured at 1024 x 1024 pixel resolution, equating to a physical size of 620 x 620 *µ*m and the z planes were spaced 2 *µ*m apart. Images were collected using a BP624/40 filter and with two-photon excitation using a MaiTai MP laser (Spectra Physics) tuned to 880nm. A second channel with a BP440/20 filter was recorded to image the dura via second harmonics. This approach ensured that the equivalent cranial depth was recorded in each experiment. Baseline images were taken for all regions of interest (ROIs). Animals were then intravenously injected with 5 *µ*g of CXCL4 diluted in saline or vehicle control (saline alone). Two or three ROIs were acquired for each animal and three or four animals were used per group.

##### Analysis

Maximum-intensity projection images of the first 60 *µ*m of tissue including the dura were generated and the fluorescence intensities (mode) for each ROI was calculated. These values were taken for each image at 120 minutes following CXCL4/vehicle injection and plotted as fold change from the baseline maximum-intensity projection images.

#### Analysis of chemokine binding to cell surface GAGs

This analysis was adapted from a previous approach (*17*). Briefly, 1 × 10^5^ genetically engineered CHO GS-/- or HEK293 6e cells were harvested from suspension culture and washed in 1 x PBS before being incubated with biotinylated recombinant human CXCL4 or CCL2 (Protein Foundry LLC) at 10 *µ*g/mL in 1 x PBS with 1% FBS for 30 min at 4 °C. Cells were washed with 1 x PBS with 1% FBS and incubated with 1:2000 Alexa Fluor 488-streptavidin (S32354, Invitrogen) in 1 x PBS with 1% FBS for 30 min at 4 °C. After washing with 1 x PBS with 1% FBS, cells were resuspended in 1 x PBS with 1% FBS and fluorescence intensity was analysed on a SA3800 spectral cell analyzer (SONY). For the heparin inhibition assays, 10 *µ*g/mL CXCL4 or CCL2 was pre-incubated with varying concentrations of porcine mucosal heparin in 1 x PBS with 1% FBS for 30 min at RT prior to incubation with cells. All experiments were performed a minimum of three times using triplicate samples.

##### Biophysical chemokine:GAG interaction analysis (Bio-layer interferometry)

Bio-layer interferometry was performed on an Octet Red96 system (Sartorius AG, Goettingen, Germany) using a methodology adapted from (*10*). Firstly, GAGs were biotinylated at their reducing end using an approach described elsewhere (*44*) and immobilised onto High Precision Streptavidin (SAX) biosensors (Sartorius AG, Goettingen, Germany). To immobilise, SAX biosensors were hydrated for 10 minutes in BLI buffer (10 mM Hepes, 150 mM NaCl, 3 mM EDTA, 0.05% Tween-20, pH 7.4). For the heparin dp8 GAG immobilisation was undertaken in BLI buffer at 0.078 *µ*g/ml to achieve an immobilisation level of approx. 1 nm. For the differentially de-sulphated heparin fractions immobilisation was undertaken at 5 *µ*g/ml in BLI buffer until all surfaces were saturated. Sensors were then washed in regeneration buffer (0.1 M Glycine, 1 M NaCl, 0.1% Tween, pH 9.5) and re-equilibrated in assay buffer. Blank reference or GAG coated sensors were then submerged into wells of black-walled 96 well plates containing 200 *µ*L of BLI buffer containing chemokines at the indicated concentrations for the indicated time (association) before being transferred to assay buffer containing wells (dissociation) before finally being washed in regeneration buffer. The binding signal was recorded throughout and signal from binding of chemokine to blank (no immobilised GAG) sensors and by GAG immobilised sensors in assay buffer alone was subtracted from that produced by chemokine binding to GAG immobilised sensors. As well as signal over time the maximum signal during the association phase of the interaction was recorded and is plotted. Data were acquired at 5 Hz and analysed using the Octet HT 10.0 analysis programme. For inhibition analysis signal was plotted against inhibitor concentration and fitted for IC_50_ values using non-linear regression in the Prism software package (GraphPad).

#### Analytical ultra-centrifugation

CXCL4 and CXCL4 K50E were re-suspended in water to a concentration of 1mg/ml and further diluted 1/10 in PBS. Samples were loaded into 2-sector cells with PBS as a reference and centrifuged at 50,000 rpm in a 4-hole An60Ti rotor monitoring the absorbance at 230 nm until sedimentation was reached. The time-resolved sedimenting boundaries were analysed using Sedfit (*45*). The resulting profiles are shown in Gussi (*46*).

##### Fluorescence recovery after photo-bleaching (FRAP) of films of in-plane mobile GAGs

Heparan sulphate used in FRAP assays was biotinylated at the reducing end (HS-b), as described previously (*44*). HS was sourced from porcine intestinal mucosa, had a molecular weight of 12 kDa, a polydispersity of 1.6 and an average of 1.4 sulfates per disaccharide (*47*). Dioleoylphosphatidylcholine (DOPC) and dioleoylphosphatidylethanolamine-CAP-biotin (DOPE-CAP-b) lipids were purchased from Avanti Polar Lipids (Alabaster, AL, USA). Streptavidin was labelled with Atto 565 fluorophores (Sav Atto 565), and all chemicals for buffers were supplied by Sigma. Working buffer used for all experimental steps consisted of 10 mM HEPES, pH 7.4, and 150 mM NaCl in ultrapure water.

##### Preparation of model glycocalyces (films of in-plane mobile GAGs)

Films of one-end anchored heparan sulfate polysaccharides on supported lipid bilayers were formed as previously described (*48, 49*). Briefly, pre-conditioned glass coverslips (24 × 24 mm^2^, type 1.5; Menzel Gläser, Braunschweig, Germany) were attached, using a bi-component glue (Picodent, Wipperfürth, Germany), to a custom-built teflon holder, thus forming the bottom of four identical wells with a volume of 50 μL each. Supported lipid bilayers (SLBs) were formed by the method of vesicle spreading (*50*), through exposure of the glass surface to small unilamellar vesicles (100 μg/mL for 30 min) made from DOPE-CAP-b in a DOPC background (molar ratio 0.5:99.5). SAv Atto 565 (40 μg/mL for 20 min) and HS-b (10 μg/mL for 20 min) were then sequentially applied to the SLBs, to anchor HS via SAv to the biotin-presenting SLB. After each incubation step, the surface was washed with a working buffer to remove excess molecules from the solution phase.

##### Fluorescence recovery after photobleaching (FRAP) assays

With HS-b anchored to SAv atto 565, FRAP of the fluorescently labelled SAv was used as a reporter of GAG in-plane mobility. Fluorescence images were taken with a confocal laser scanning microscope (Zeiss LSM880, Zeiss, Germany) using ZEN software. The objective was a Plan-Apochromat 40×/1.4 Oil DIC M27, and the light source a 561 nm laser. The pinhole size was 198 μm (5 Airy units). In each FRAP experiment, 3 pre-bleach images of the surface (177.12 × 177.12 μm^2^; 256 × 256 pixels) were acquired before a circle in the centre of the image (10 μm radius) was bleached, and 57 images were acquired post bleach to monitor the fluorescence recovery.

To quantify fluorescence recovery, all images were analysed in Fiji (*51*). Mean fluorescence intensities were extracted from two regions of equal radius (6.92 μm) in each image in the sequence: the bleach region was located at the image centre and the reference region at the periphery. Fluorescence intensities were corrected for background fluorescence and intensity fluctuations, and normalised using the following equation: 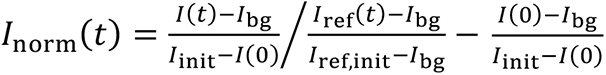. Here, *I*_bg_ is the background intensity (measured in the absence of fluorophores, and constant across the image), *I*_init._and *I*(*t*) are the intensities in the bleach region prior to bleaching (*t* < 0) and at post-bleach time *t* (*I*(0) = *I*(*t* = 0)), respectively, and *I*_ref,init._ and *I*_ref_(*t*) are the corresponding intensities in the reference region. With this normalisation, the unbleached (or fully recovered) intensity is 1, and the intensity in the bleach region immediately after bleaching is 0. Recovery curves were plotted as normalised intensity against time, and the area under each curve from *t* = 0 to *t* = 40 s was calculated to represent an effective measure of the in-plane mobility of SAv (and the attached HS) on the SLB.

To assess the impact of CXCL4 on HS mobility, the HS film was incubated with CXCL4 alone (250 nM), with a mix of CXCL4 and RTO (at 2:1 molar ratio), or with a mix of CXCL4 and a control IgG (at 2:1 molar ratio) for 1.5 h; the HS film was then analysed by FRAP (within 10 minutes, and with excess proteins in the solution phase). Prior to use, the mixtures of CXCL4 and RTO (or control IgG) were incubated for 1 h to allow complexes to form. Three independent experiments were performed for each condition, and in each experiment three FRAP series were acquired at different positions on the surface. Data shown in Fig. 4E and Suppl. Fig. 7 represent the mean ± 95% confidence intervals across all 9 data sets per condition.

#### Immgen

The Immgen data base was probed using the MyGeneSet data browser (*14*).

#### Statistics

All statistics were performed using the Prism (GraphPad) software package. Experiments containing two groups were analysed using an unpaired T-test and data containing more than two groups were analysed using a one-way ANOVA with a post-hoc multiple comparison test. p < 0.05 was considered to be statistically significant.

## Acknowledgements

The authors would like to acknowledge numerous colleagues for reading and helping in drafting the manuscript. The Bioimaging Facility microscopes used in this study were purchased with grants from BBSRC, Wellcome Trust and the University of Manchester Strategic Fund. Special thanks to Peter March, Roger Meadows and Steven Marsden for their help with the microscopy. FRAP analyses was performed on a microscope within the Bioimaging Facilities at University of Leeds, supported by the Wellcome Trust (WT104818MA).

## Funding

DPD was supported by a Sir Henry Dale fellowship jointly funded by the Wellcome Trust and Royal Society (Grant Number 218570/Z/19/Z) and a Wellcome Trust centre grant (Grant Number R125761). RLM was supported by The Carlsberg Foundation CF20-0412. JT and RLM were supported by The European Union’s Horizon 2020 Research and Innovation Programme 899687, HS-SEQ.

## Author contributions

Conceptualization: ALG, RLM, DPD

Methodology: ALG, RK, ARER, AJLR, NP, CH, LMR, HLB, CLS, EAY, RPR, JET, TMH, GJG, TAJ, IS, RPR, DPD

Investigation: ALG, RK, ARER, AJLR, NP, CH, LMR, HLB, CLS, EAY, RPR, TAJ, IS, DPD

Funding acquisition: RLM, DPD

Project administration: DPD

Supervision: TMH, GJG, RPR, RLM, DPD

Writing – original draft: RLM, DPD

## Competing interests

The authors declare no competing interests

## Data and materials availability

Data and materials are available on request.

## Notes

### Competing Interest Statement

The authors have declared no competing interest.

