## Supplementary figures for "Chemokine CXCL4 interactions with extracellular matrix proteoglycans mediate wide-spread non-receptor mediated immune cell recruitment"

**Supplementary Figure 1.**

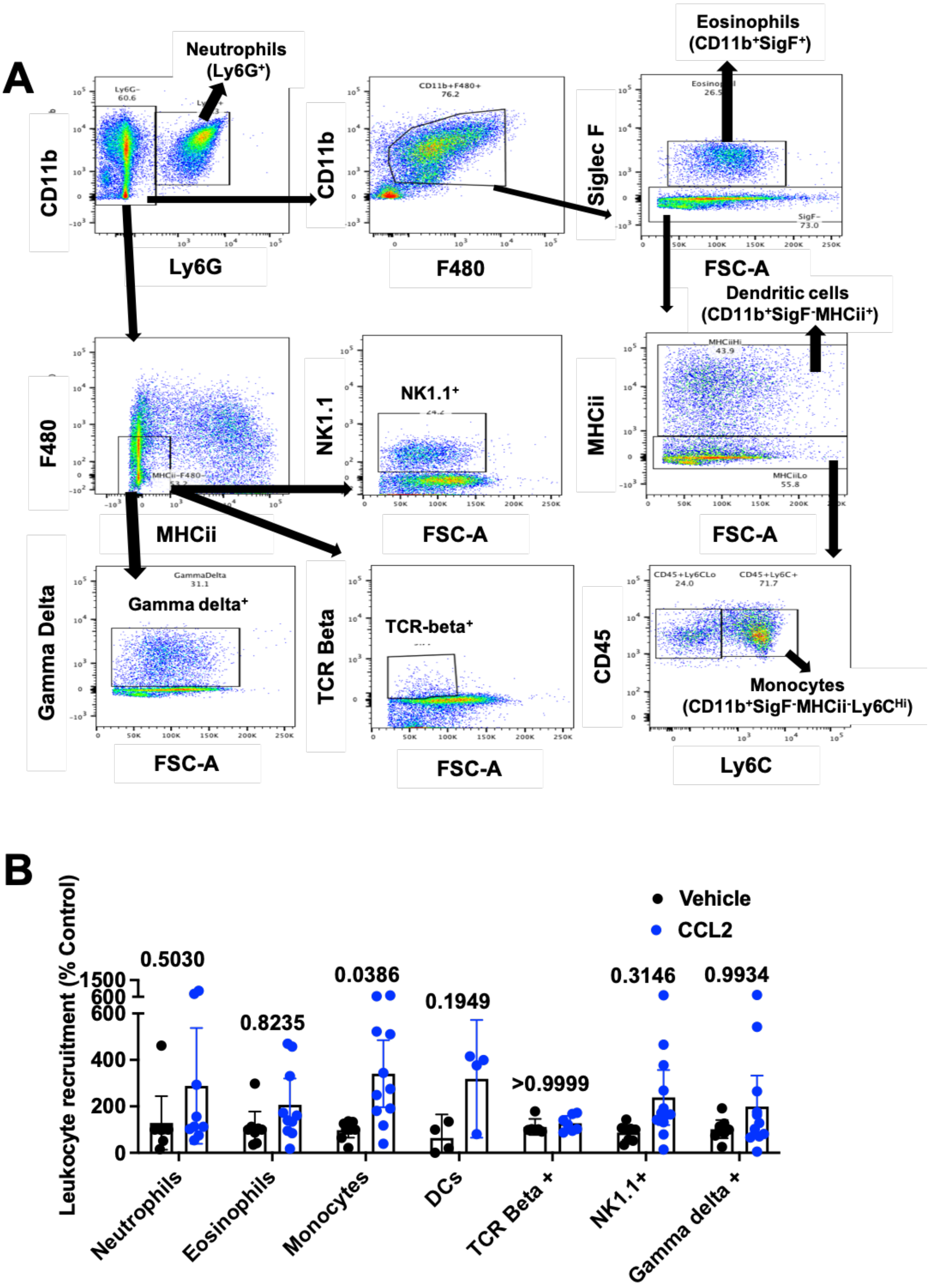

**Supplementary Figure 1. Analysis of chemokine mediated recruitment of leukocytes to the murine air pouch.** (A) Gating strategy used for flow cytometry analysis of cells retrieved from the air pouch cavity. (B) Quantification of CCL2 mediated leukocyte recruitment to the air pouch.

B is the mean with 95% confidence intervals and represents at least two separate experiments where data have been pooled and each dot represents an individual mouse. Data in B is normalised relative to vehicle controls. Individual p values are shown and was analysed using a one-way ANOVA with a post-hoc Sidak analysis of log-transformed data.

**Supplementary Figure 2.**

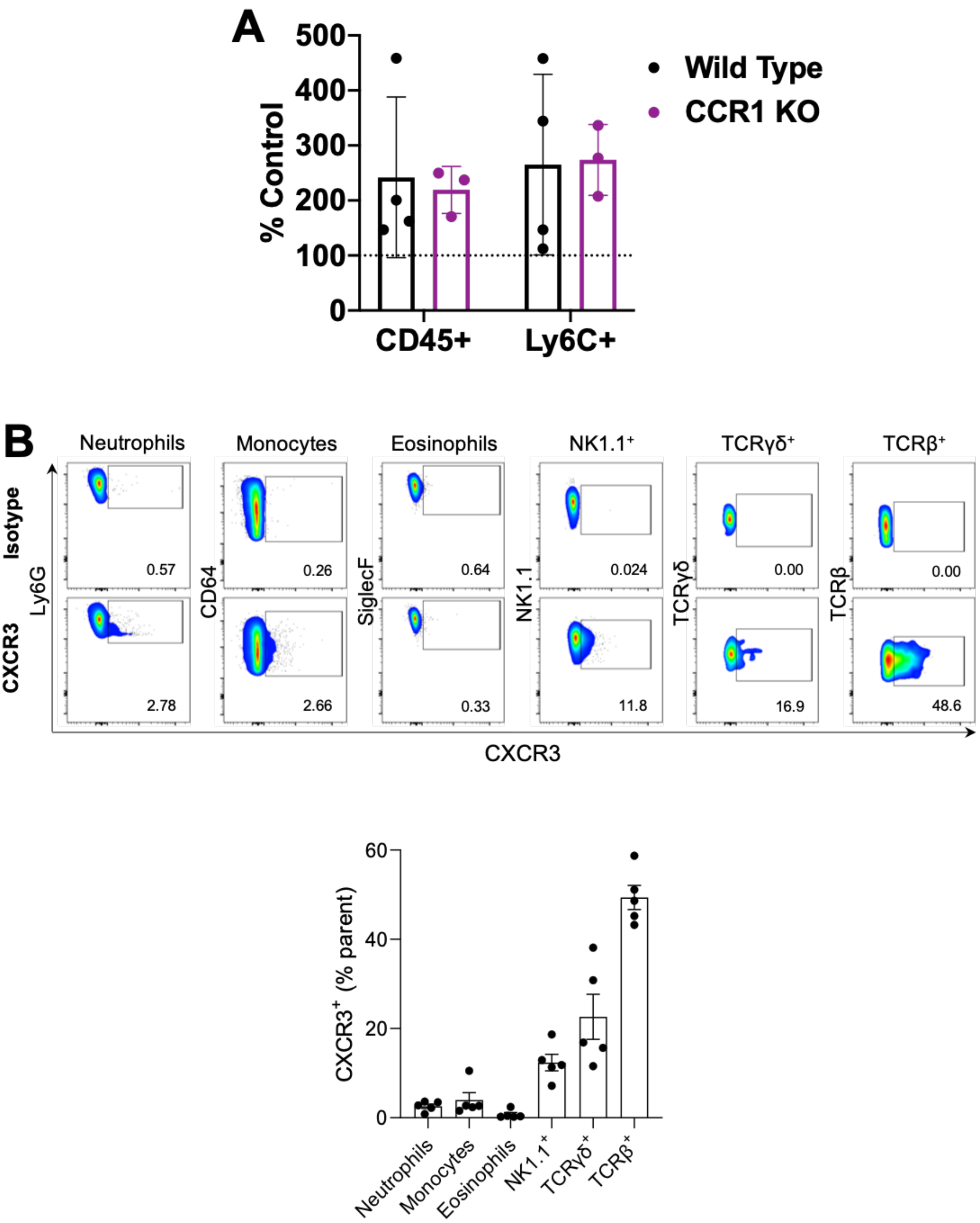

**Supplementary Figure 2. CXCL4 mediated recruitment is independent of CCR1 and CXCR3.** (A) CXCL4 (5 μg) was injected into dorsal air pouches in wild type and CCR1 KO mice and 24 hours later the recruited cells were analysed using flow cytometry. (B) Indicated cell types were analysed for CXCR3 expression using flow cytometry, squares (gates) are drawn on flow cytometry plots to indicate where cells with positive staining for the leukocyte marker and CXCR3 would sit.

A and B (lower panel) are mean with 95% confidence intervals and represent at least two separate experiments where data have been pooled and each dot represents an individual mouse. Data in A are normalised relative to vehicle controls.

**Supplementary Figure 3.**

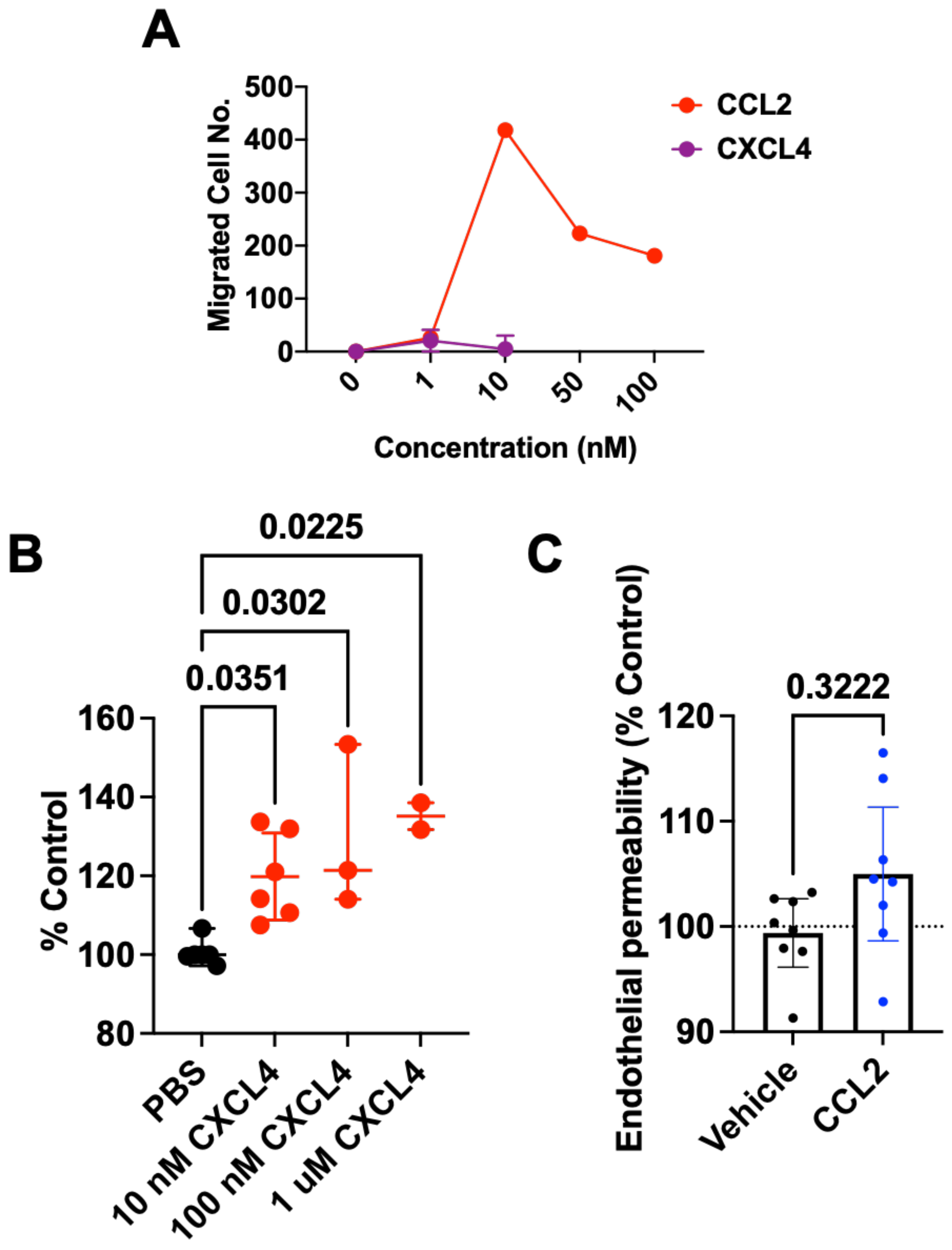

**Supplementary Figure 3. CXCL4 but not CCL2 increases endothelial permeability *in vitro*.** (A) Chemotaxis assay of human monocytes to CXCL4 or CCL2. (B) Endothelial permeability in response to vehicle control or different amounts of CXCL4. (C) Endothelial permeability with and without CCL2 (10 nM).

B and C are mean with 95% confidence intervals and represent at least two separate experiments where data have been pooled. Data in B and C is normalised relative to vehicle controls. Individual p values are shown, C was analysed using an unpaired t-test and B was analysed using a one-way ANOVA with a post-hoc Sidak analysis.

**Supplementary Figure 4.**

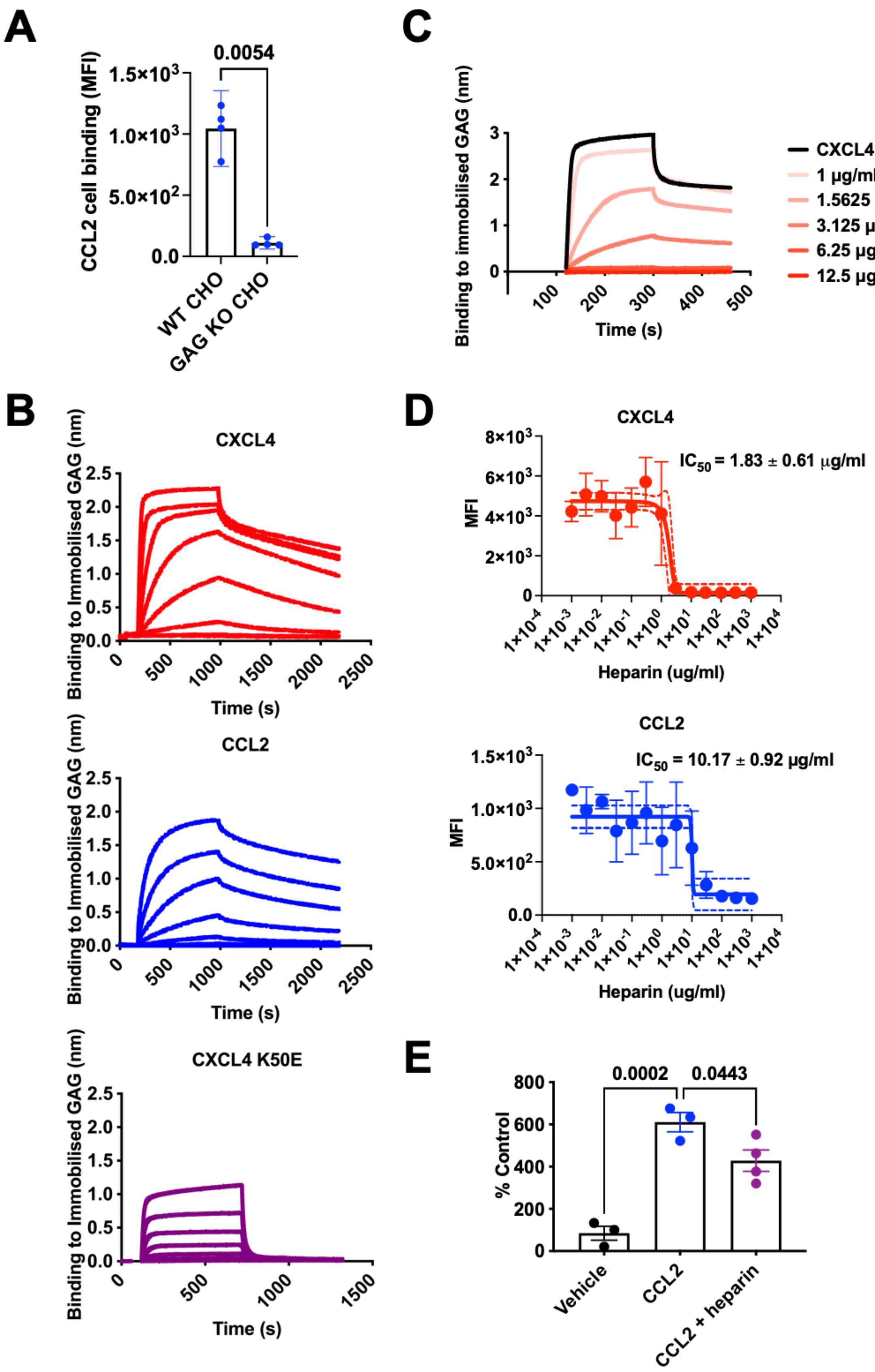

**Supplementary Figure 4. Exogenous GAG inhibits CXCL4 and CCL2 binding to the cell surface.** (A) CCL2 binding to CHO cells with and without surface GAGs (B4Galt7 KO). (B) CXCL4 (500, 250, 125, 62.5, 31.3, 15.6, 7.8 or 3.9 nM), CCL2 (1000, 500, 250, 125, 62.5, 31.3, 15.6 or 7.8 nM) or CXCL4 K50E (1000, 500, 250, 125, 62.5 or 31.3 nM) were passed over a heparin dp8 surface followed by buffer alone to analyse the interaction over time using BLI. (C) CXCL4 binding to immobilised heparin dp8 over time on its own and with varying concentrations of competing GAG (heparin) in the solution phase. (D) The binding of labelled chemokine to CHO cells was monitored with and without pre-incubation with different concentrations of exogenous GAG (heparin) and analysed for $IC_{50}$ values. Solid line represents fit of the data using non-linear regression and dotted lines represent 95% confidence intervals of the fit. (E) Analysis of leukocyte (Ly6$C^{Hi}$ monocyte) recruitment to the air pouch in response to CCL2 with and without exogenous heparin.

A and E are mean with 95% confidence intervals and all plots represent at least two separate experiments. Data in E is normalised relative to vehicle controls. Individual p values are shown, A was analysed using an unpaired t-test and E was analysed using a one-way ANOVA with a post-hoc Dunnett analysis.

**Supplementary Figure 5.**

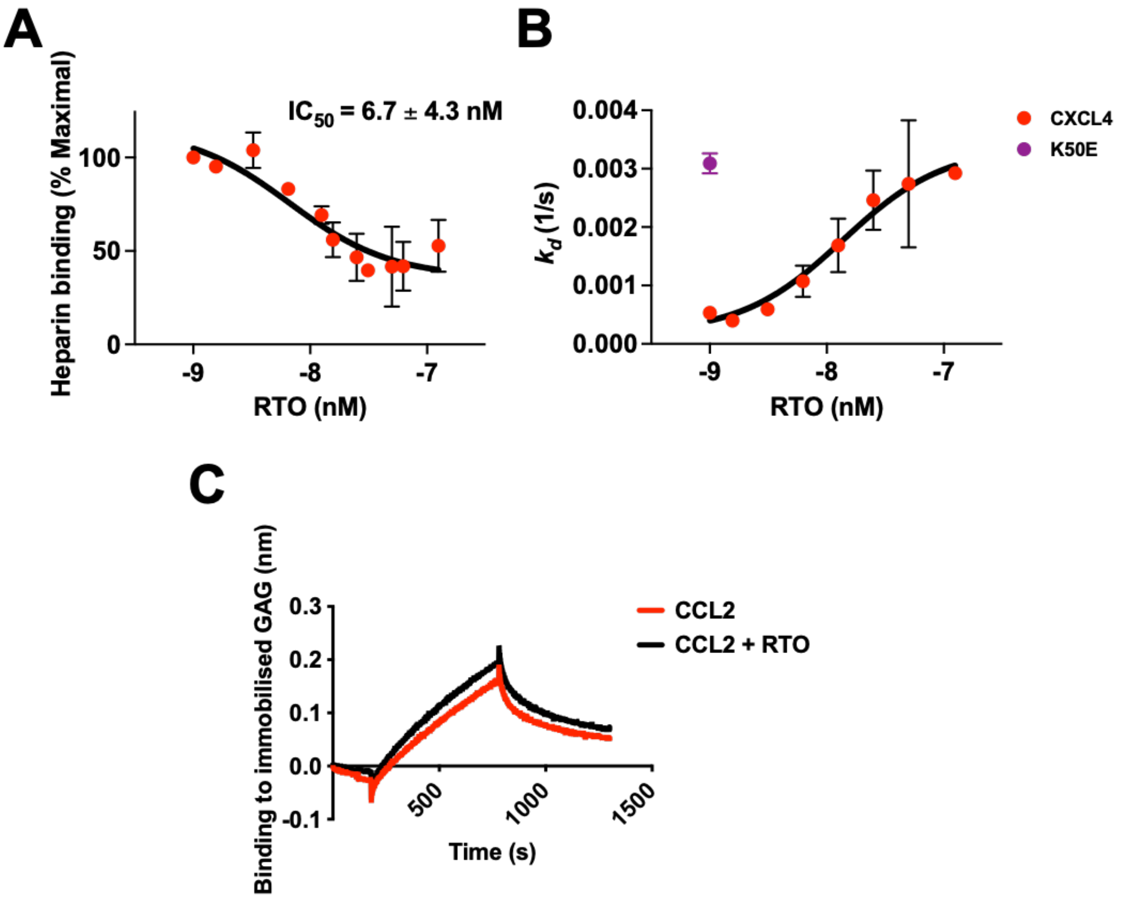

**Supplementary Figure 5. CXCL4 oligomerisation drives its' GAG interaction.** (A) BLI assay final CXCL4 signal (after washing) is plotted with and without pre-incubation with RTO antibody. (B) Using BLI the CXCL4:dp8 off-rate ($k_d$) was determined and plotted with and without RTO antibody; CXCL4 mutant K50E off-rate included for comparison. (C) CCL2 (50 nM) with or without the anti-CXCL4 antibody RTO (50 nM) was passed over a heparin dp8 coated sensor and the subsequent interaction was analysed.

All plots represent at least two separate experiments. (A and B) solid line represents fit of the data points using non-linear regression analysis.

**Supplemental figure 6.**

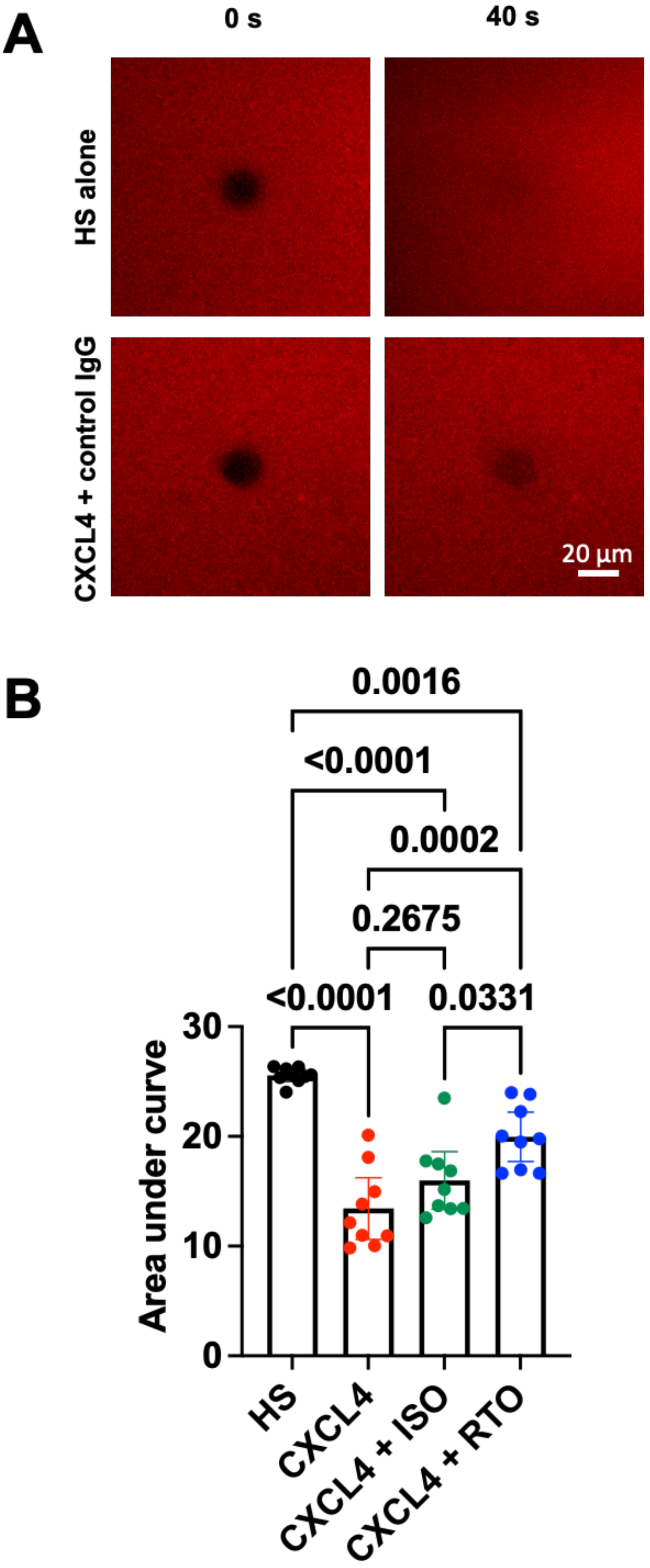

**Supplementary Figure 6. FRAP analysis of CXCL4 mediated GAG cross-linking.** (A) Images of the bleached region with HS alone or CXCL4 isotype control immediately after bleaching (0s) or 40s after bleaching. (B) Fluorescence recovery over time analysed through the areas under the curves in Fig. 4I (between 0 and 40 s). Each dot represents a separate experiments per condition, and is the mean from three recovery curves taken at different positions on the surface. B was analysed using a one-way ANOVA with a post-hoc Dunnett analysis.

**Supplemental figure 7.**

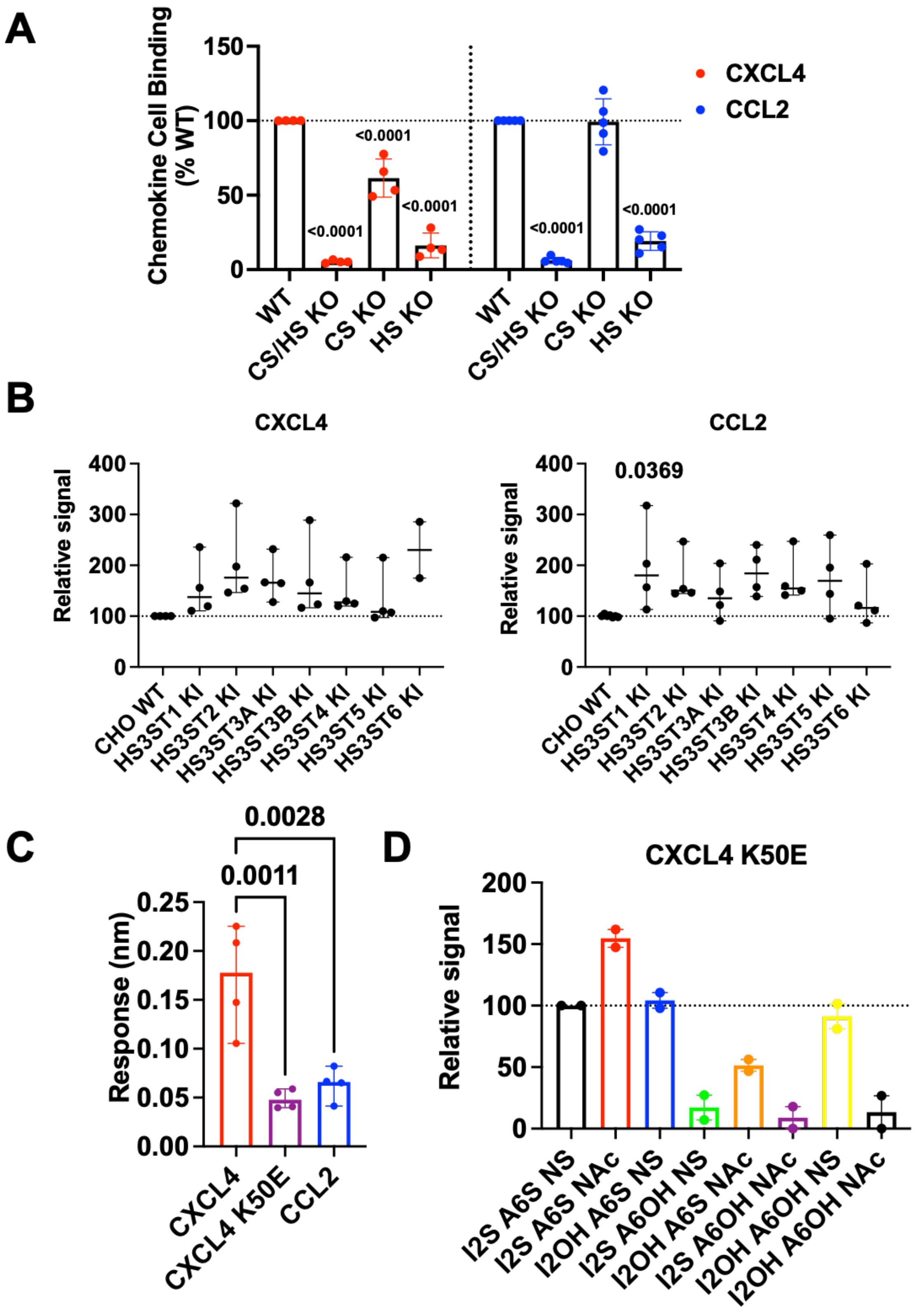

**Supplementary Figure 7. GAG fine structure mediates selective binding and cell surface retention of chemokines.** (A) Chemokine binding to HEK cells with no, HS alone, CS alone or HS and CS combined GAGs knocked out. (B) Chemokine binding to CHO cells with 3-O GAG sulphation enzymes knocked in. (C) Maximum BLI signal of chemokine (500 nM) binding to fully sulphated heparin (2-O, 6-O and N- sulphated). (D) BLI maximum signal of CXCL4 K50E (500 nM) binding to saturated surfaces of the indicated fully- or de- sulphated heparin samples.

Plots in A-D are mean with 95% confidence intervals and represent at least two separate experiments. Data in A, B and D are normalised relative to wild type cells (A and B) or fully sulphated heparin fragments (D). Individual p values are shown and A, B and C were analysed using a one-way ANOVA with a post-hoc Sidak analysis. p values in A and B are comparisons of the indicated groups to WT controls.
